# A computational framework for detecting signatures of accelerated somatic evolution in cancer genomes

**DOI:** 10.1101/177261

**Authors:** Kyle S. Smith, Debashis Ghosh, Katherine S. Pollard, Subhajyoti De

## Abstract

By accumulation of somatic mutations, cancer genomes evolve, diverging away from the genome of the host. It remains unclear to what extent somatic evolutionary divergence is comparable across different regions of the cancer genome versus concentrated in specific genomic elements. We present a novel computational framework, SASE-mapper, to identify genomic regions that show signatures of accelerated somatic evolution (SASE) in a subset of samples in a cohort, marked by accumulation of an excess of somatic mutations compared to that expected based on local, context-aware background mutation rates in the cancer genomes. Analyzing tumor whole genome sequencing data for 365 samples from 5 cohorts we detect recurrent SASE at a genome-wide scale. The SASEs were enriched for genomic elements associated with active chromatin, and regulatory regions of several known cancer genes had SASE in multiple cohorts. Regions with SASE carried specific mutagenic signatures and often co-localized within the 3D nuclear space suggesting their common basis. A subset of SASEs was frequently associated with regulatory changes in key cancer pathways and also poor clinical outcome. While the SASE-associated mutations were not necessarily recurrent at base-pair resolution, the SASEs recurrently targeted same functional regions, with similar consequences. It is likely that regulatory redundancy and plasticity promote prevalence of SASE-like patterns in the cancer genomes.

## INTRODUCTION

A saturation analysis indicated that most of the common, recurrent drivers in protein coding regions in all major cancer types are likely to be detected by the genome projects (Lawrence *et al*., 2014; International Cancer Genome Consortium *et al*., 2010; Collins and Barker, 2007). In contrast, our understanding of cancer-associated mutations in non-coding regions, which cover ∼98% of the genome is so far preliminary at best. Most of the fixed germ line mutations and single nucleotide polymorphisms associated with complex traits and diseases are noncoding, and it is suspected that non-coding regulatory mutations may have important roles in tumorgenesis as well (Khurana *et al*., 2016) – providing a rationale for tumor whole-genome sequencing. While so far only a relatively small number of recurrent noncoding mutations with regulatory functions have been identified, emerging findings suggest that there might be additional novel regulatory changes active in cancer genomes (Chang *et al*., 2016; Scott W Piraino and Furney, 2017; Imielinski *et al*., 2017).

While a major emphasis of the ongoing efforts focus on detection of recurrent noncoding mutations, we adopt a different, evolution-driven approach. Locus-specific accelerated evolution in the human lineage (Human accelerated regions) marked by accumulation of a significant excess of germ line mutations has been associated with functional regulatory changes in loci regulating the development of the central nervous system, limb and other organs linked to human-specific traits(Pollard *et al*., 2006). With accumulation of somatic mutations, cancer genomes also evolve(Podlaha *et al*., 2012), diverging away from the genome of benign, progenitor cell lineages in the host. Somatic mutation frequency varies between genomic regions, such that the extent of evolutionary divergence varies considerably between different regions. We present a novel computational framework, called SASE-mapper, to identify genomic regions that show signatures of accelerated somatic evolution (SASE) in a subset of samples in a cohort, and used it to scan 365 cancer genomes from 5 cohorts representing 4 different cancer types to identify signatures of accelerated somatic evolution at a genome-wide scale.

## METHODS

### Definition of SASE

Acquired somatic alterations indicate how much a genomic region within a tumor genome has evolved from the germline sequence of the host(Podlaha *et al*., 2012). We defined a signature of accelerated somatic evolution (SASE, **Figure 1A**), which is marked by a significant excess of somatic mutations compared to that expected based on the local background mutation frequency, typically in a subset of the samples in a cohort. Therefore, it would be distinct from the typical mutational hotspot signatures. We argue that loci carrying SASE could be potential targets of context-directed mutagenesis and/or selection during tumor evolution and should be investigated for functional and clinical significance.

**Fig. 1.**
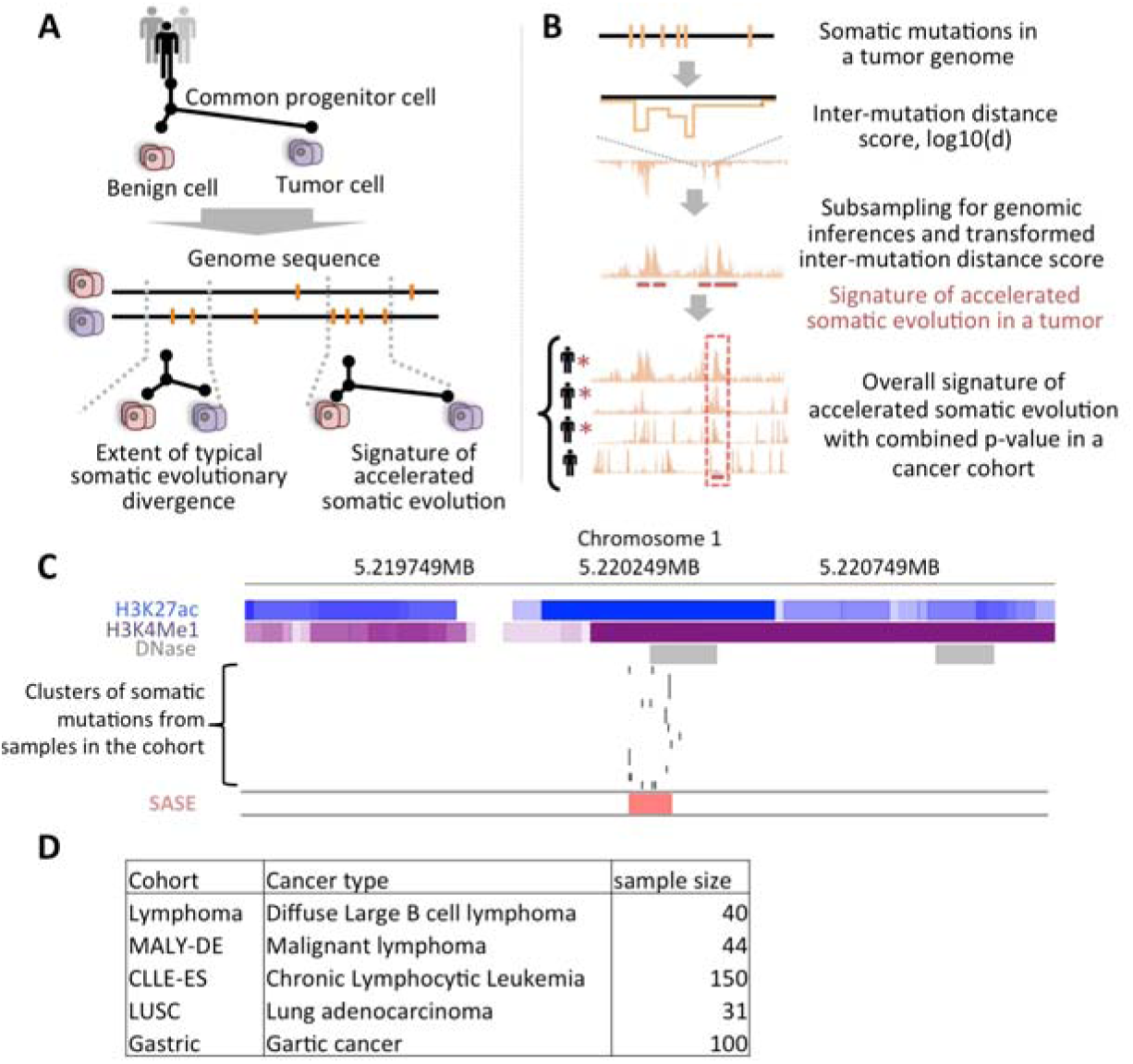
(A) Signature of accelerated somatic evolution, (B) Schematic representation showing SASE-mapper analysis pipeline. (C) Annotated GASTRIC-SASES region from gastric cancer in predicted enhancer. (D) Summary of the datasets used in the study.

### Equivariant genomic regions

While identifying SASE, we need to consider that local, background mutation frequency varies considerably between different genomic regions(Lawrence *et al*., 2014) in a context dependent manner. Factors such as evolutionary conservation, chromatin, GC content, and replication timing play major roles in modulating local mutation rate. Therefore, we binned the genome into 10bp blocks for which the mean GERP, PhastCons, and phyloP evolutionary conservation scores were used to classify the region as under high (default: GERP>=1.5, phastCons>=0.4, phyloP>=1), low (default: GERP<=0, PhastCons<=0.1, phyloP<=0) or medium conservation. Similarly, we classified genomic regions as early, late, or variable replication timing regions using published approach(Pedersen *et al*., 2013). We used similar approaches to classify genomic regions according to their GC content (low<=20%, medium<=60%, high> 60%) and giemsa-staining based chromatin (euchromatin, intermediate, heterochromatin) status(Speir *et al*., 2016).

We segmented the genome based on the conservation (low, medium, high), replication timing (early, late, variable), GC content (low, medium, high), and chromatin status of the region (euchromatin, intermediate, heterochromatin,), and defined regions as equivariant if they had equivalent status for these major covariates of mutation rate. Mutation frequency had low variation within EVRs but high variation between EVR classes (**Supplementary Fig. 1**), indicating that our approach accommodates major regional variations in mutation frequency. We note that other covariates could be incorporated while defining the EVRs, but in doing so the number of segments increases and the total size of the genomic segments assigned to an EVR class decreases, such that there may not be enough somatic mutations within certain EVR classes for meaningful downstream analyses. Similarly, selecting only equivalent tri-nucleotide context within respective EVRs for subsampling is possible, but in some cases it reduces the number of possible sites for permutation to generate a meaningful null distribution. Therefore, we used a set of EVRs characterized by the combination of conservation and replication timing, which showed similar results, but was considerably faster for downstream analysis.

### Detection of signatures of accelerated somatic evolution

Let there be N number of completely sequenced cancer genomes in a cohort, each having n_i_ (i∈N 1, 2…N) somatic mutations. For each sample, inter-mutation distances between successive mutations were calculated (**Figure 1B**), and the corresponding probability to detect such inter-mutation distances in the tumor genome was estimated. For this purpose, we generated a context-aware null distribution of inter-mutation distances using a variation of the subsampling approach(Bickel *et al*., 2010) by randomly shuffling mutations among respective equivariant regions within the chromosome in that sample. This preserved the context-dependent differences in background mutation density and biases arising from known covariates in the null model, and allowed us to estimate the probability of detecting certain inter-mutation distances by chance alone. Genomic regions with clusters of mutations would have peaks of –log10(*p*-values) while the valleys should mark regions where mutations are sparse. Importantly, the transformation removes biases due to overall somatic mutation burden, or common regional covariates(Lawrence *et al*., 2013), allowing us to compare the landscape of *p*-values between different genomic regions in a cancer genome, between cancer genomes, or even between samples across cancer types that have different background burden of somatic mutations. It also allowed us to combine signals from adjacent regions in the genome, as described below.

By our definition, a biologically relevant peak should be present in a number of samples, though it need not be present in all samples in the cohort. Therefore, we combined p-values from all samples in the cohort using the truncated product method(Zaykin *et al*., 2002), prioritizing the candidate SASE regions that show p-values with a chosen threshold (default: <=0.05 in >=3 samples). Final SASE regions were determined by identifying peaks in -log10 p-values through comparison of the expected values per region to those observed by means of the significant fold change (SFC) metric(Knijnenburg *et al*., 2014). SFC combines the p-value and effect size to determine enrichment and can be interpreted as the lower bound on the effect size. At a predefined significance level (default: p-value < 1e-6) SFC is the effect size that is significant. Adjacent genomic segments with p-values above the p-value threshold and within a specified distance (default: 30bp) were combined, and the contiguous genomic segments and their mean significant fold-changes were reported.

### Dataset

Genome-wide somatic mutation data for 365 samples from 5 different cancer cohorts(International Cancer Genome Consortium *et al*., 2010; Collins and Barker, 2007; Berger *et al*., 2012; Morin *et al*., 2013; Wang *et al*., 2014) (**Figure 1C**) was obtained, and the somatic mutations were mapped to hg19. Each cohort had at least 30 samples. Typically the samples in the cohorts were sequenced using Illumina sequencing technology at ≥30X. While different cohorts might have processed samples differently and called mutations using their preferred pipeline, observed concordance of SASE signatures in related cancer types indicates that our results are robust against biases due to variant calling and batch effects.

### Genomic context and functional elements

We obtained data for DNase hypersensitivity and ChIP-seq based transcription factor binding site data for human cell lines (Consortium, 2012), Ensembl predicted enhancers (Segway and chromhmm predicted), vistaEnhancers(Visel *et al*., 2007), and EnhancerFinder step1(Erwin *et al*., 2014) predicted developmental enhancers. As an example, we show in gastric cancer the representative region with SASE shown in **Figure 1C** overlapped with predicted DNase hypersensitive open chromatin, H3K4Me1 marks and H3K27ac marks. Functional element enrichment was determined by comparing identified SASE regions and the flanking region by way of a one-sided binomial test. 3D nuclear context and long-range interaction patterns of the SASE regions were evaluated based on data for lamin-associated domains (Guelen *et al*., 2008) and HiC based chromosome conformation (Rao *et al*., 2014), respectively. HiC data binned at 100kb resolution was downloaded from Rao et al.(Rao *et al*., 2014) for 5 cell lines. Regions with more than 3 reads, normalized by the method previously proposed by (Lieberman-Aiden *et al*., 2009), and mapped with at least a 30 mapping quality score in at least 3 of the 5 cell lines were considered. Signatures of accelerated somatic evolution co-occurring in 3 samples in addition to supporting hic data were determined to be interacting. The GM12878 cell line data was used for lymphoma and chronic lymphocyte leukemia cohorts. We obtained genomic context tier annotations from a previous report(Mardis *et al*., 2009).

### Expression and clinical data analysis

A subset of the cohorts analyzed (Collins and Barker, 2007) also had paired gene expression and clinical data available. When a SASE was detected in the promoter of a gene, its expression level (normalized read counts) was compared between the groups of samples that had the signature and other samples in the cohort using Mann-Whitney U test. Survival differences of the samples between the groups were examined using log-rank statistic and Kaplan Meier plot. All p-values were corrected for multiple testing using Benjamini-Hochberg false discovery rate.

### Implementation

SASE-mapper is implemented in Python 2.7. It can run on ∼2 million somatic mutations from 25 samples on a machine with 4 processors and 8GB of RAM in less than 10 minutes. Cohorts with larger number of samples and higher mutation burden may take longer. The rate-limiting step is the estimation of combined p-values. A user can change default values for most of the parameters. Relaxed parameter settings can permit the detection of broad SASEs, while stringent parameters tend to yield narrower regions.

## RESULTS

### Validation analysis

We evaluated SASE-mapper predictions by a three-step validation and comparative assessment to examine whether it can detect realistic examples, before applying it to the cohorts of completely sequenced cancer genomes (**Figure 1D**). First, we generated a cohort with 30 synthetic tumor genomes with background somatic mutation frequency of ∼10/Mb, and spiked 10% of the samples (n=3) with an excess of mutations (3-5 per sample) at 5 arbitrarily chosen regions (length 50bp to 2000bp, **Figure 2A**, **Supplementary Fig. 1**). Default parameters were determined to detect all 5 of the spike-ins. We repeated the analysis with synthetic cohorts with different background mutation rates, cohort sizes, and proportion of samples carrying the spike, and obtained similar results (**Figure 2B**, **Supplementary Fig. 1**).

**Fig. 2.**
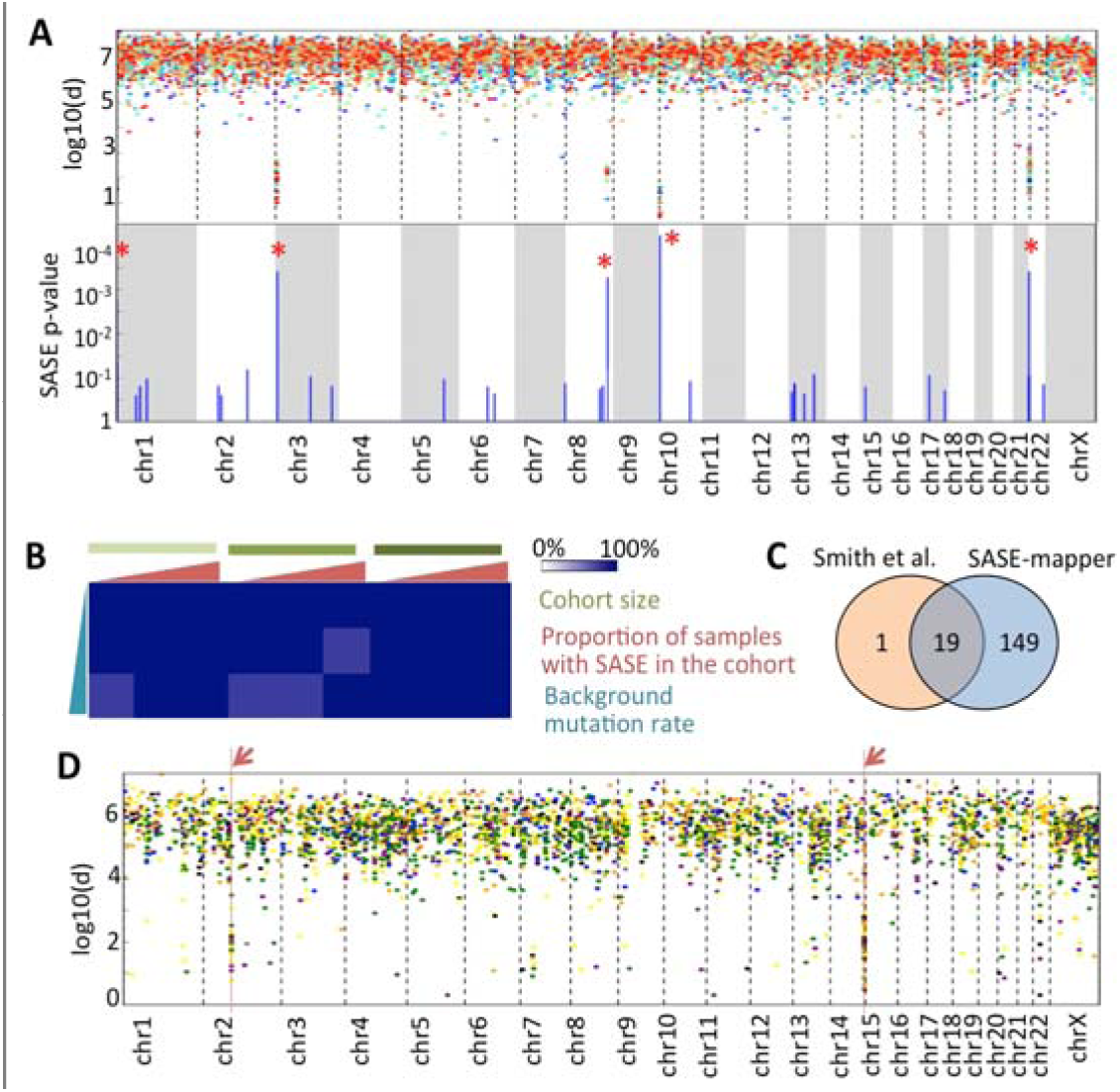
(A) Upper-panel shows raindrop plot of Inter-mutation distances, *d*, of 30 simulated samples ol which 5 were spiked with hyper mutation at predetermined loci, while the lower panel shows the SASE p-values. Loci with spikes are marked with asterisks. All the spiked regions were correctly identified. (B) The analysis was repeated with synthetic cohorts with different background mutation rates, cohort sizes, and proportion of samples carrying the spike. In 85% cases all spiked regions were identified, and in 100% cases 280% spiked regions were identified. (C) Venn Diagram showing the overlap between SASE-mapper and Smith et al. in detecting SASE in gene promoter regions, (D) Raindrop plot of inter-mutation distances in a CIL sample with kataegis like regions indicated by arrow.

Second, using a different, promoter-specific algorithm we previously identified evidence for accelerated somatic evolution in promoters of known cancer genes such as MYC and BCL2 in lymphoma, which were associated with altered expression and poor clinical outcome(Smith *et al*., 2015). Of the 20 regions detected in that promoter-centric analysis, 19 were also detected by our current, unbiased genome-wide analysis (**Figure 2C**). We found an additional 149 regions with SASE elsewhere in the tumor genomes in the cohort. SASE in selected cancer gene promoters, including those detected in this analysis, were associated with altered expression of the cancer gene and clinical outcome(Smith *et al*., 2015). This suggests that our genome-wide approach can identify known and novel target regions. Importantly, SASE-associated changes (increase or decrease) in expression of these genes were consistent with their role in promoting tumor growth, which might be suggestive of selection during somatic evolution.

Lastly, kataegis is a pattern of localized hypermutation identified in some cancer genomes, often resulting from multiple cytosine deaminations catalyzed by AICDA/APOBEC family enzymes(Lada *et al*., 2012). In a cohort of lymphoma and chronic lymphocytic leukemia, we detected the kataegis signatures using a published definition(Lawrence *et al*., 2013), and then applied SASE-mapper to detect signatures of accelerated evolution. SASE-mapper, despite receiving no prior information about kataegis signatures, re-identified all the regions with kataegis **(Figure 2D)**. This demonstrates that our framework is able to detect biologically and clinically important instances of accelerated somatic evolution, including novel mutation signatures, in a cohort of cancer genomes.

### Signatures of accelerated somatic evolution in cancer cohorts

We analyzed somatic mutation data for 14 cohorts, composed of 1,595 samples, representing 10 different cancer types (International Cancer Genome Consortium *et al*., 2010; Collins and Barker, 2007; Berger *et al*., 2012; Morin *et al*., 2013; Wang *et al*., 2014) using SASE-mapper to detect signatures of accelerated somatic evolution at a genome-wide scale. SASE-mapper used equivariant regions defined based on evolutionary conservation and replication timing (**Supplementary Fig. 2**), which captured most of the regional variation in background mutation rate, but an extended model incorporating chromatin, GC content, and other factors yielded similar results (**Methods**). Overall, we detected 4,759 SASE regions, with the number varying across cancer types (134-3,005 per cohort; **Supplementary Table 1**). In the MALY-DE lymphoma cohort, a number of loci, including promoters of known cancer genes such as MYC and BCL2 exhibit signatures of accelerated somatic evolution (**Figure 3A; Supplementary Table 1**). To explore the reproducibility of the SASE, we compared results across independent lymphoma cohorts. Of the 168 SASE detected in either cohort, 59 were found in both cohorts. Most of these SASEs were also detected in the CLL cohort (**Figure 3B**), and these three-way hits are largely in regulatory regions of known cancer genes (**Supplementary Fig. 3**). We note that, the cohorts had different sequencing coverage and used different variant detection pipelines, which could add to technical variability. Furthermore, SASEs were typically present in a minority of the samples in a cohort, such that a moderate size cohort may not have sufficient number of samples with a given signature by chance alone. Nonetheless, the observed concordance between the CLL and two lymphoma cohorts was highly statistically significant (permutation test; p-values < 10^-10^), suggesting that the identified loci are most probably genuine targets. We also detected SASE near known cancer-associated genes in other cohorts such as lung cancer (LUSC) cohorts (**Figure 3C**, **Supplementary Table 1)**. The cohorts that most frequently share recurrent SASE are both lymphoma cohorts and CLL (**Supplementary Fig. 4**), which were also most closely related in terms of the disease etiology. Our findings are consistent with that reported by other studies. For instance, an investigation into mutational hotspots by Piraino et al.(Scott W. Piraino and Furney, 2017) identified both the MYC and BCL2 promoters to be recurrently mutated in addition to the MIR142 locus in the ICGC lymphoma cohort (International Cancer Genome Consortium *et al*., 2010). However, their method bins the genome into 50bp segments and is therefore likely to miss some other mutational hotspots of different sizes.

**Figure 3:**
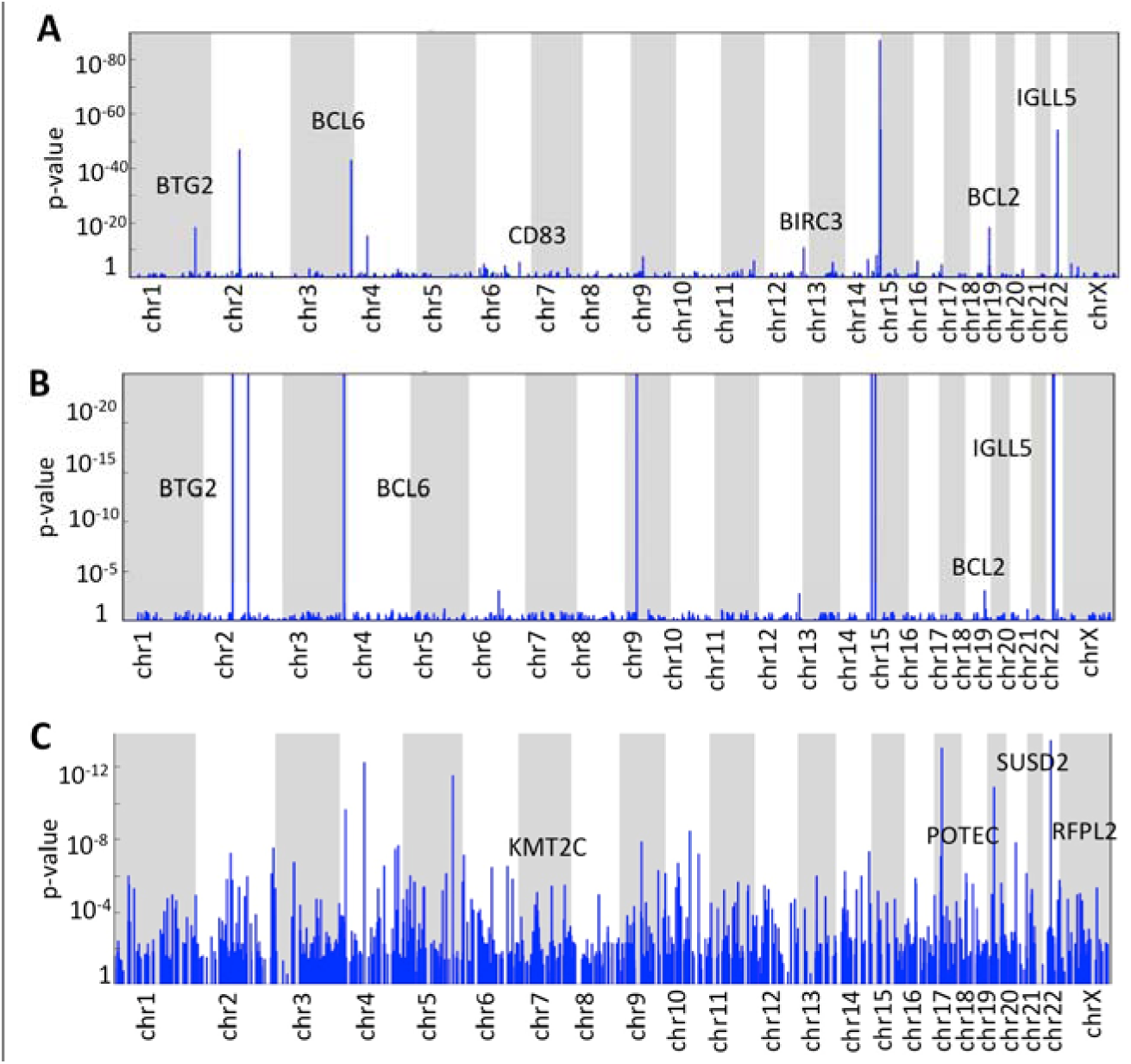
Manhattan plot showing genome-wide signatures of accelerated somatic evolution in (A) MALY-DE, (B) CLL, and (C) LUSC cohorts. Some of the SASE overlapped with promoters of known cancer genes, consistent with that found in a promoter-centric investigation previously.

**Fig. 4.**
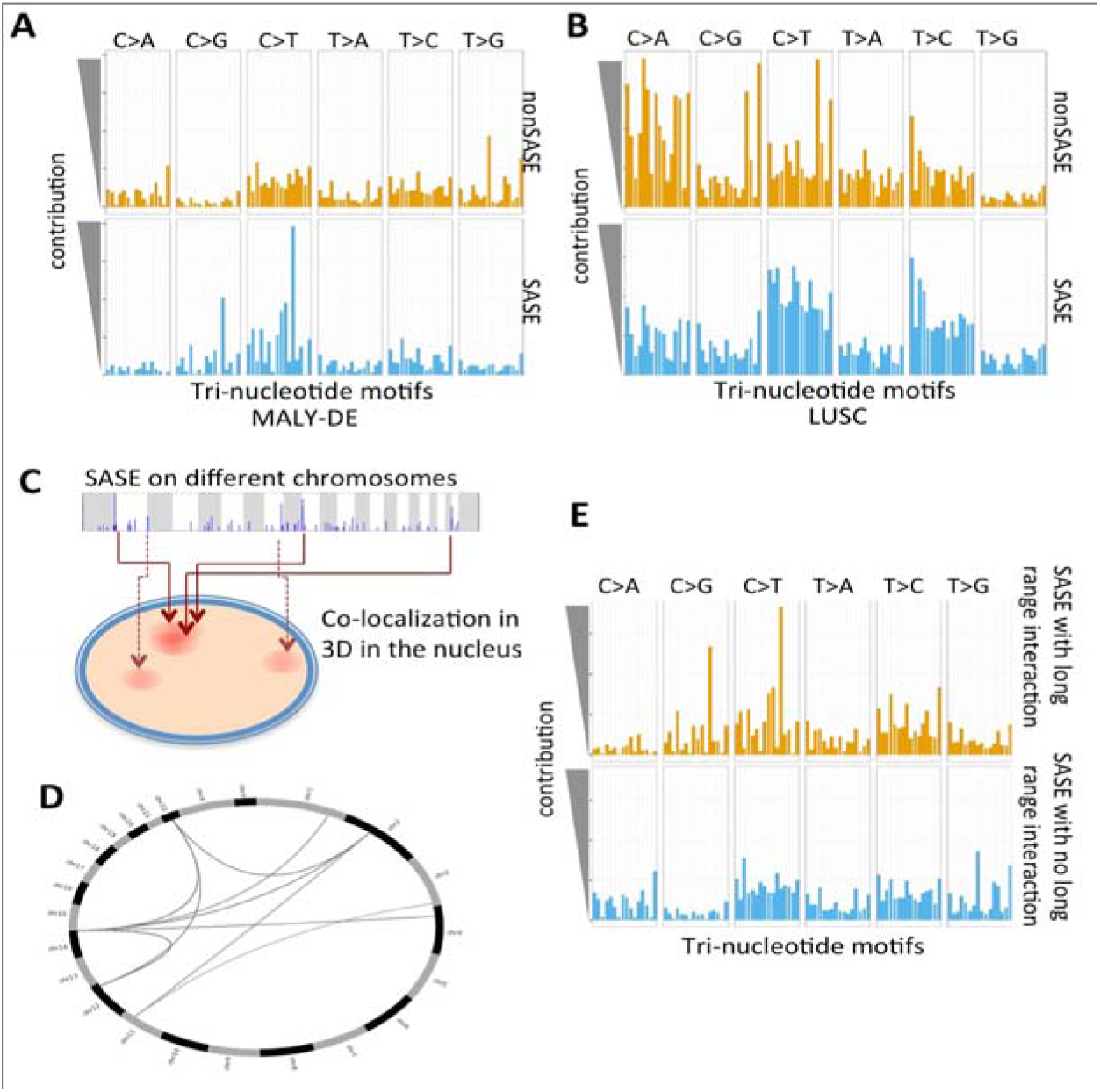
(A) Somatic mutational signatures for the MALY-DE cohort, (B)) Somatic mutational signatures for the LUSC cohort, (C) Schematic representation showing nuclear co-localization would lead to long range interactions among loci affected by SASE, (D) SASE-SASE long range interactions detected using HiC data within a lymphoma, MALY-DE, and CLLE-ES combined cohort. (E) Somatic mutational signatures for the loci affected by SASE have evidence for long range interactions in multiple samples and those that do not have such evidence in the for lymphoma, MALY-DE, and CLLE-ES combined cohort.

### Genomic context of SASE

SASE can be found on all human chromosomes and in a variety of genomic contexts, but some contexts were more common that others (**Figure 4A**). Less than 3 percent of SASEs were observed in protein-coding genes, and except for the lymphoma cohorts, coding regions were generally depleted for SASE. However, SASEs in highly conserved elements in noncoding regions were relatively more common; 36 percent of SASEs occurred in highly conserved elements throughout the genome, a vast majority of them outside coding regions. Repetitive regions showed no systematic enrichment, but it is challenging to map reads and call mutations in those regions. SASEs occurred significantly more often in the promoter regions than that expected by chance (Permutation p-values <0.009 in both lymphoma cases); as shown in **Figure 2C** and **3A-B**, we detected SASE in the promoters of known cancer genes such as BCL2, IGLL5, BTG2, and BCL6 in the lymphoma and CLL cohorts. There were additional examples of SASE in promoter regions of known cancer genes in other cohorts. For instance, SASE in regulatory regions of RNF212, a ring finger protein involved in meiotic recombination(Kong *et al*., 2008) in the lung cancer cohort was associated with altered expression of the gene product and also reduced survival of the patients carrying the signature. Transcription factor binding sites and enhancers were also significantly enriched for SASE (Permutation enrichment p-values <0.05) in selected cancer types. In general, many SASE were in open chromatin in cancer cells or had epigenomic signatures associated with activeregulatory regions, H3K4Me3 (12 cohorts have enrichment p-values <=0.05, **Figure 4A**). In gastric cancer, for example, the representative region with SASE shown in **Figure 1C** overlapped with predicted DNase hypersensitive open chromatin, H3K4Me1 marks, and H3K27ac marks, and had clusters of mutations in 14 of the 100 samples. SASEs frequently overlapped with Ensembl(Consortium, 2012)-predicted binding sites of CTCF (36% of cohorts have p-values <=0.05, **Figure 4A**), which play important roles in long range chromatin looping. It is possible that some of these mutations might alter the long-range chromatin interactions, but it needs to be further investigated conclusively.

### Some regions with SASE cluster together in the 3D in the nucleus

We revisited the SASEs in the context of 3D nuclear organization of genomic DNA integrating data on lamina associated domains (Guelen *et al*., 2008) and HiC-based long-range interactions (Rao *et al*., 2014), outlined in the **Methods**. SASEs were more frequently associated with consistently inter-lamina regions (iLAD) that are at the nuclear interior compared to the consistently lamina-associated regions (cLAD) in the nuclear periphery (**Figure 4A**). For instance, of the 168 detected SASE in MALY-DE, 19 occur in cLAD and 95 occur in iLAD (Permutation test; p-values <=0.05). Similar patterns were also observed in all other cohorts. We analyzed the landscape of somatic mutations in the lymphoma cohort in their 3D nuclear context (**Figure 4B**) by integrating data on nuclear localization and chromosome conformation capture (HiC)-based long range interaction data for lymphoblastoid cell lines(Rao *et al*., 2014), which would be developmentally related to the progenitor cell of origin of lymphoma. Nearly 25 percent of SASEs in the MALY-DE cohort had at least minimal evidence for interaction (≥ 1 predicted HiC interaction, see **Methods**) with another SASE (detected in the same sample), the majority of which were previously identified APOBEC targets **(Figure 4C, Supplemental Fig. 5)**. The SASEs present in a majority of the cancer gene promoters in the lymphoma and CLL cohorts had evidence for SASE-SASE long-range interactions, indicating their likely shared origin. Another interesting example was the signatures of accelerated somatic evolution on chromosome 17 at MIR142 locus, a known AICDA target(Robbiani *et al*., 2009), which had evidence of interaction with 10 other SASEs in the same samples. Similar SASE-SASE long-range interactions were detected in both the lymphoma cohorts and CLL cohort **(Figure 4C)**, indicating that such overlaps in the 3D are unlikely due to chance alone. Evidence for long-range SASE-SASE interactions was also identified in about 15 percent of signatures present in the melanoma and lung squamous cell carcinoma cohort **(Supplemental Fig. 5)**. Signatures of accelerated somatic evolution on chromosome 1 between 142.5Mb – 142.7Mb, detected recurrently in LUSC and gastric. Thus, many SASE regions mapped to different regions within and across chromosomes could be spatially clustered in the 3D in the nucleus, and might arise from the same mutagenic process.

**Figure 5:**
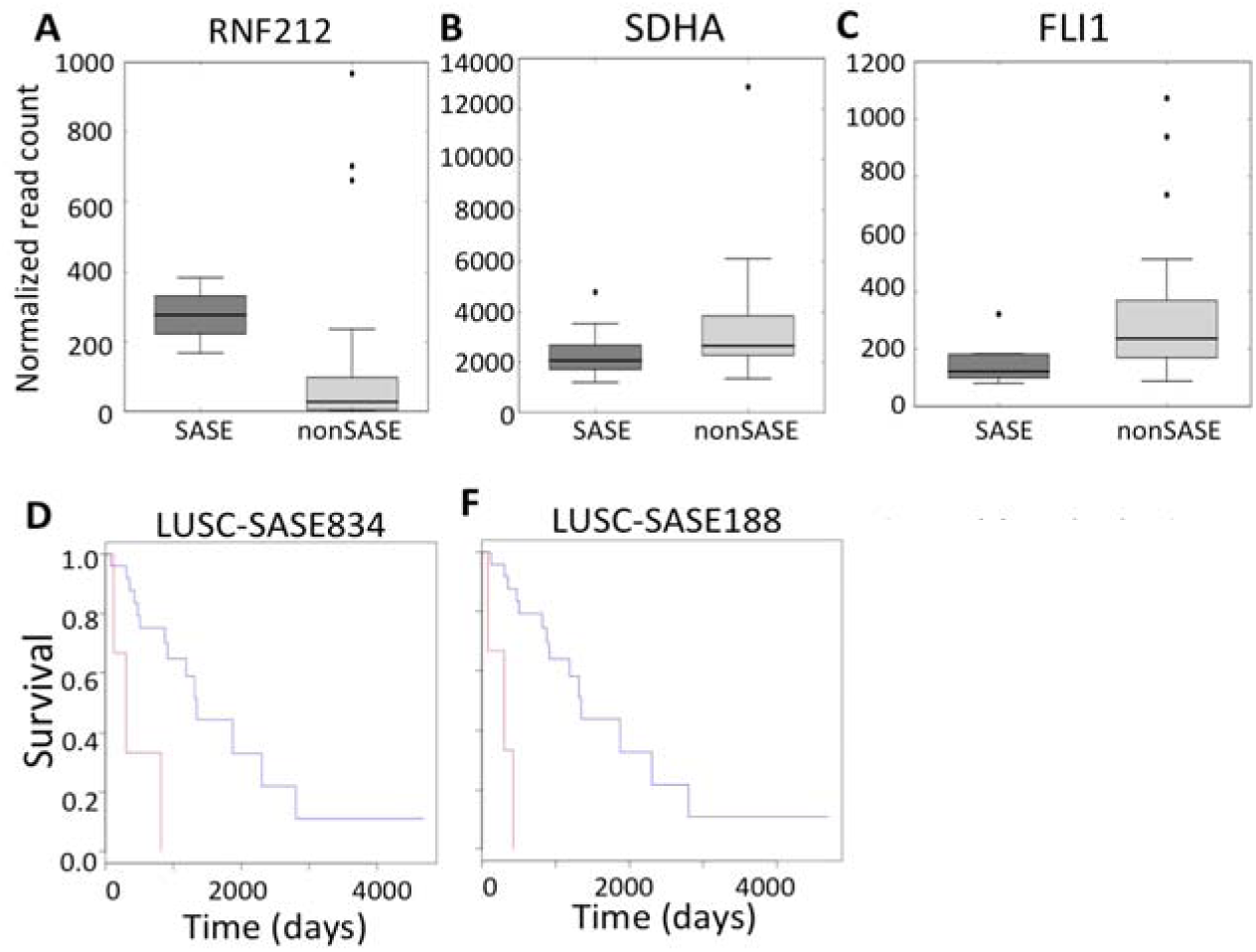
(A) Boxplot showing mRNA expression difference for RNF212 between the samples that have signatures of accelerated somatic evolution and other samples in the lung cancer cohort. (B) and (C) show the similar results for SDHA and Fill respectively in the LUSC cohort. Kaplain-Meyer Plots show survival analysis for selected SASE regions in the LUSC cohort (D-F).

### SASE associated mutation signatures

We used non-negative matrix factorization-based method (Alexandrov *et al*., 2013) to infer the likely basis of mutation signatures that characterize SASE. This analysis indicated that APOBEC/AICDA associated mutagenesis is frequent in SASEs, especially in the lymphoma and CLL cohorts (**Figure 4A-B** and **Supplementary Fig. 6**). In both the lymphoma cohorts, SASEs exhibit increased instances of C> T and C> G mutations compared to frequencies observed in non-SASEs, which is suggestive of AICDA/APOBEC mediated hyper-mutation(Alexandrov *et al*., 2013; Lada *et al*., 2012) of these regions (**Figure 4A,** p-values < 10^-10^, Chi Squared test). Perhaps unsurprisingly, many of these affected genes with SASE in the promoters were also known targets of AICDA. In the CLL samples, T> C substitutions, and C> G or C> T at the G<u>C</u>T context were proportionally more common in SASE regions (p-value < 0.05, Chi Square test). T> C is a classic signature of APOBEC cytidine deaminase mutagenesis pattern, but the mechanistic basis for the substitution preference at G<u>C</u>T context is unclear. Interestingly, the SASEs that had mutational signatures consistent with AICDA activity also had strong evidence for SASE-SASE long-range interactions **(Figure 4D)**. It is likely that AICDA activity in the transcription factories, where genomic loci that are distant on the linear DNA but interact in the 3D nuclear territory during transcriptional regulation, contributed to the observed patterns. In the lung cancer cohort, C> A substitutions, which are frequently associated with smoking, were relatively depleted in SASE-affected regions, and instead transversions (C> T and T> C substitutions) were relatively enriched (**Figure 4B**; p-value < 10^-^10^^, Chi Square test). SASE-affected regions in gastric cancer were also enriched for C> T and T> N mutations. Finally, SASE frequently overlapped with predicted kataegis-like signature in different cancer cohorts (**Supplementary Fig. 7**).

### Regulatory and clinical significance of SASE

Genomic and nuclear context of the SASE-associated regions led us to investigate whether SASE, especially those proximal to the genes, are associated with regulatory alterations (**Figure 5**; **Supplementary Table 1**). In the malignant lymphoma cohort, SASE-mapper detected SASE in 45 regions (median mutations ≥2; present in ≥3 samples) that occurred in predicted regulatory regions (FDR corrected p-values <= 0.05, see **Methods**). Of those, 40 showed altered expression in genes within 2MB (cis-interactions) and 5 were associated with alterations in age. For instance, known cancer genes such as MYC, BCL2, IGLL5, etc. had altered expression (Smith, Yadav et al. 2015). In the lung squamous cell carcinoma cohort, SASE-mapper detected evidence for accelerated somatic evolution in 155 regions (median mutations ≥2; present in ≥3 samples) that overlapped with predicted enhancers (FDR corrected p-value <= 0.05). Of those, 105 showed altered expression in genes within 2MB. For instance, about 25 percent of samples contained mutations in a SDHA promoter, which was associated with decreased mRNA expression of the gene. A SASE present at telomeric region of chromosome 4 (chr4p16.3) was associated with alterations in expression of 28 genes in a 2MB region including cancer-related genes such as FGFRL1, SH3BP2, CRIPAK, and POLN. Similarly, SASE on chromosomes 7p11.2 and 11q25 were associated with increased expression of PSPH, and decreased expression of FLI1, respectively. SASE present in a RNF212 intron was associated with altered expression in samples with mutations distributed throughout the locus (**Figure 5A-C**).

In the lung squamous cell carcinoma cohort, we detected SASE an exon of KMT2C, which was significantly associated with reduced survival. KMT2C (MLL3) is a lysine methyltransferase that is frequently mutated in different types of cancers(Song *et al*., 2014; Herz *et al*., 2014). SASE present in TAF4 and FLT4 were also associated with reduced survival. In addition, signatures of accelerated somatic evolution in intron of RNF212(Kong *et al*., 2008), which correlated with altered mRNA expression, was associated with reduced survival as well. Of the 105 SASEs occurring in predicted, non-coding regulatory regions, 4 are associated with reduced survival (**Figure 5D-F**).

## DISCUSSION

We suspect that the SASE probably arises due to a combination of both mutagenesis and selection. In a subset of cases, context specific mutagenesis plays a key role (e.g. kataegis and APOBEC signatures in lymphoma). Positive selection or relaxation of purifying selection also appears to be important in some other cases, especially those associated with deregulation of cancer gene expression and also aggressive tumor progression leading to poor survival. It remains unclear whether all the mutated sites within a SASE in a tumor are under positive selection, especially since they are not typically recurrent at the base-pair resolution in multiple samples in a cohort. One possibility could be that, inherent regulatory redundancy in noncoding regions enables different mutations or their combinations to achieve similar regulatory consequences, a concept gaining traction recently. An alternate, but not necessarily mutually exclusive, possibility is that context specific mutagenesis generates an excess of mutations at a given locus, majority of which are passengers that hitchhike one or a few driver mutations. Furthermore, some mutations might directly affect oncogenic pathways by creating or perturbing transcription factor binding sites, while others could have more subtle effects e.g. modulation of nucleosome occupancy, changes in local chromatin or DNA conformations such that their epistatic interactions ultimately change the regulatory environment. Further work needs to be done going beyond association to establish causality. In any case, when evidence for recurrent noncoding regulatory mutations is sparse, our findings make a case for assessment of non-traditional mutational signatures in non-coding regions of cancer genomes.

## ACKNOWLEDGEMENTS

KS and SD conceived the project. KS performed the experiments. KS, DG, KSP, and SD analyzed the data and wrote the manuscript. Funding: T15 LM009451 [to KS], Lung Cancer Research Foundation, Boettecher Foundation [to S.D.]; Grone Endowed Chair of the University of Colorado Cancer Center [to DG]; Gladstone Institutes, a gift from the San Simeon Fund, and P01 HL089707 [to K.S.P.];

## SOFTWARE AVAILABILITY

SASE-mapper is written in Python 2.7. The source-code, manual, and example files are available at http://github.com/kylessmith/SASE-mapper.

